# PolyMorph3D: A flexible MATLAB-based tool for quantifying microglia morphology and multi-colocalization in three dimensions

**DOI:** 10.64898/2026.11.19.767700

**Authors:** Andrew R. Yoblinski, Thien Quy Pham, Jyoti Dubey, Marianne L. Seney

**Affiliations:** Department of Psychiatry, University of Pittsburgh School of Medicine, Pittsburgh, PA 15213; Center for Neuroscience at University of Pittsburgh, Pittsburgh, PA, 15261; Department of Biostatistics and Health Data Science, University of Pittsburgh, Pittsburgh, PA, 15261; Department of Bioengineering, University of Pittsburgh, Pittsburgh, PA, 15261

## Abstract

As the principle immune cell of the central nervous system (CNS), microglia have been extensively studied throughout development and in a wide range of disease states. Of special focus have been their morphological features such as branching complexity, physical cell-cell interactions with surroundings, and gene expression readouts relating to phagocytosis and cytokine signaling. Each of these domains offers a window onto microglia function, and high-resolution confocal microscopy has long been a workhorse technique to visualize all three. A comprehensive view of microglia function requires both high-fidelity quantification with 3D resolution and colocalization of microglia with functional proteins such as those related to phagocytosis and with nearby structures such as synapses. Accordingly, a number of analytical tools have been developed to measure these features. However, existing open-source options lack either colocalization or 3D capability, while commercial solutions can be costly. Here, we sought to develop an open-source solution that addresses each of these limitations. PolyMorph3D is a MATLAB-based program capable of yielding individual cell-level data of microglia morphology and overlap with up to 4 other sources, all in 3D. The pipeline can be used effectively in mouse tissue and in human postmortem brain, thereby offering a flexible, low cost, and high-fidelity solution for the analysis of microglial morphology and function.

## INTRODUCTION

Microglia are brain-resident macrophages, yet are morphologically dissimilar to macrophages in other tissues, resembling instead the ramified morphology of their neighbors (e.g., astrocytes) in the parenchyma. Under homeostatic conditions, microglia extend and retract processes to survey the local environment. Should a disturbance be detected, microglia take on new morphological phenotypes to respond to disrupted brain homeostasis (Davalos et al., 2005). Leaning on microglia morphologies observed during nerve injury or neurodegeneration, several approaches were developed to categorize microglia as disease associated (DAMs) (Keren-Shaul et al., 2017), by adopting classical macrophage nomenclature of M1 vs M2 (Michelucci et al., 2009), or along a functional continuum (e.g., ramified, primed, reactive, ameboid) (Torres-Platas et al., 2014; Wu et al., 1992). However, researchers are increasingly cognizant of limitations that categorization places on evaluation of microglia and their functional states (Paolicelli et al., 2022). The field is shifting towards continuous readouts of microglia morphology and function. Interrogating microglia across multiple modalities is particularly important, as single measurements (e.g., branching complexity) do not always correlate with functional outputs (e.g., phagocytic activity) (Yoblinski et al., 2025).

Several tools can quantify morphological features of microglia. The ideal solution would have five features: 1) high-throughput, 2) high-fidelity for 3-dimensional structure, 3) individual cell-level resolution, 4) colocalization with other channels, and 5) accessible (i.e., low/no cost; prior coding experience not required). While these features are present among existing tools, no one tool exhibits all five (Table 1). The nearest options are the commercially available programs Imaris (Oxford Instruments) and Neurolucida*®* 360 (MBF Bioscience). Each is high-fidelity, includes individual cell-level resolution, and offers channel colocalization. However, both can be labor intensive, and licenses prohibitively expensive, making them inaccessible for many. Several published low-cost options use MATLAB (Heindl et al., 2018; Salamanca et al., 2018; York et al., 2018) or ImageJ/FIJI (Benkeder et al., 2025; Hansen et al., 2022; Kugler et al., 2023) (ImageJ/FIJI is free; MATLAB license is comparatively inexpensive and sometimes available to academic researchers at no cost). However, these open-source options lack the capability to colocalize microglia with signal from other channels, missing out on indicators of microglia function.

**Table 1.**

| Tool | Platform | High Throughput | High Fidelity in 3D | Single Cell Resolution | Colocalization | Accessible |
|---|---|---|---|---|---|---|
| PolyMorph3D | MATLAB | ✓ | ✓ | ✓ | ✓ | ✓ |
| 3DMorph (York et. al. 2018) | MATLAB | ✓ | ✓ | ✓ | ✕ | ✓ |
| MIC-MAC (Salamanca et. al. 2018) | MATLAB | ✓ | ✓ | ✓ | ✕ | ✓ |
| MMQT (Heindl et. al. 2018) | MATLAB | ✓ | ✓ | ✓ | ✕ | ✓ |
| MorphoCellSorter (Benkeder et. al. 2025) | ImageJ/FIJI | ✓ | ✕ | ✓ | ✕ | ✓ |
| MotiQ (Hansen et. al. 2022) | ImageJ/FIJI | ✓ | ✓ | ✓ | ✕ | ✓ |
| Neurolucida 360 (MBF Bioscience) | proprietary | ✕ | ✓ | ✓ | ✓ | ✕ |
| Imaris (Oxford Instruments) | proprietary | ~ | ✓ | ✓ | ✓ | ✕ |

To address these limitations, we developed user-friendly, GUI-based MATLAB scripts based on the published 3DMorph script (York et al., 2018), with significant changes and key additions. Our scripts preserve much of the 3DMorph workflow, starting by identifying microglia as binary volumes in 3D, but with added flexibility in thresholding and improved preview options. Our script also adds the option to pre-filter and/or merge microglia channel signal with a secondary microglia-related channel. We have added the capability to process overlap with up to 4 additional channels, with flexibility in user-defined refinement of colocalization options. Finally, we migrated and modified the “Automatic Mode” of 3DMorph to separate scripts better optimized for batch processing. Here, we demonstrate the flexibility and fidelity of our pipeline in mouse and human brain tissue. We offer a low cost, high-throughput, and high-fidelity pipeline requiring no prior coding experience for analysis of microglial morphology and function in 3D.

## MATERIALS AND METHODS

### Mouse Immunofluorescence

Tissue from male and female *Thy1-YFP* mice was used to generate test images for developing the pipeline. All mice were social housed with same-sex littermates on normal 12h/12h light/dark cycle. All animal procedures were performed in accordance with the University of Pittsburgh animal care committee’s regulations. For all mouse data presented, mice were anesthetized with isoflurane and transcardially perfused (5mL/min) with ice cold phosphate buffered saline (PBS) followed by 4% paraformaldehyde (PFA) in PBS. Brains were post-fixed in 4% PFA for at least 24 hours then cryoprotected in 30% sucrose in PBS for at least 24 hours prior to sectioning on a cryostat. Brains were cut in 40µm coronal sections and stored in glycerol-based storing solution at –20°C. Free floating sections were incubated with the following primary antibodies: rabbit anti-Iba1 (Fujifilm Wako, cat. #01919741; 1:1000) and rat anti-CD68 (BioLegend, cat. #137001; 1:1000). Corresponding fluorescent goat secondary antibodies were applied and visualized in two channels, 647 (Invitrogen, cat. #A21245; 1:1000) and 555 nm (Invitrogen, cat. #A48263; 1:1000), respectively. Images from medial prefrontal cortex were acquired on an Olympus FV1200 laser scanning confocal inverted microscope. A 60× oil immersion objective (NA 1.40) was used for all imaging. Image acquisition was performed using Olympus Fluoview software. Z-stacks were acquired at 0.17 µm intervals, and laser powers for each channel were individually optimized to avoid under- or overexposure.

### Human Postmortem Brain Tissue/Immunofluorescence

Postmortem human brain tissue was obtained during routine autopsies conducted by the Office of the Allegheny County Medical Examiner (Pittsburgh, PA), following informed consent from next-of-kin. All procedures were approved by the University of Pittsburgh’s Committee for Oversight of Research and Clinical Training Involving Decedents and the Institutional Review Board for Biomedical Research. Fixed coronal sections (40 µm) from the left hemisphere were collected across the rostro caudal extent of the subgenual anterior cingulate cortex (sgACC; Brodmann area 25). Free-floating tissue sections were incubated with the following primary antibodies: rabbit anti-Iba1 (Fujifilm Wako, cat. #01919741; 1:100), Invitrogen™ Alexa Fluor™ 568 Phalloidin (Fisher Scientific, cat. # A12380), guinea pig anti-Shank3 (Synaptic Systems, cat. #162304; 1:150), and CD68/SR-D1 Goat anti-Human, Polyclonal, R&D Systems (Fisher Scientific cat. #AF2040-SP; 1:200). Corresponding fluorescent donkey secondary antibodies were applied and visualized in four channels (720, 640, 568, and 488 nm, respectively). Image acquisition was performed using a Crest inverted microscope equipped with an Olympus spinning disk confocal unit. A 60× oil immersion objective (NA 1.40) was used for all imaging. Image acquisition and initial processing were performed using Olympus Cell Sens software. Z-stacks were acquired at 0.15 µm intervals, and exposure times for each channel were individually optimized to avoid under- or overexposure. Lipofuscin autofluorescence was captured in a dedicated channel (excitation 405 nm) using fixed exposure time across all samples to facilitate consistent post hoc removal during image processing. Raw image stacks were deconvolved in Olympus Cell Sens software to reduce out-of-focus light and improve signal resolution. Following deconvolution, Gaussian subtraction was applied in MATLAB to all channels except the microglial marker channel (Iba1), to reduce low-intensity background fluorescence.

### Neurolucida*®* 360 Analysis

Three-dimensional reconstruction and analysis of microglia were performed using Neurolucida 360. Regions of interest (ROIs) were defined as a central volume within each image stack, excluding approximately 20 µm from the lateral edges and 4 µm from both the top and bottom of the z-stack to minimize edge artifacts. Within each ROI, individual microglial cells (including soma and processes) were manually segmented and reconstructed in 3D. Using Neurolucida*®* 360’s puncta detection module, fluorescence signals were quantified based on spatial proximity to microglial structures. Specifically, Lipofuscin puncta were masked within 12 µm of microglial structures. CD68-positive puncta were identified within 0.2 µm, excluding regions overlapping with lipofuscin. Batch analysis of morphometric and puncta-related parameters was performed using Neurolucida Explorer, including quantification of soma size, branching complexity, and puncta colocalization with microglia.

### Microglia Validation Analysis

To validate our workflow, we performed two analyses. First, we selected 10 images from the human post-mortem dataset described above and analyzed them using PolyMorph3D. To validate that our post-threshold filtering steps were effective (details in Results), we compared identified objects in filtered and unfiltered images using paired t-tests for 2-group comparisons and repeated measures ANOVA (rmANOVA) followed by Bonferroni-adjusted post-hoc comparisons for 4-group comparisons. Second, we selected 40 microglia at random from the Neurolucida*®* 360 trace dataset described above. The TIFF images containing these microglia were processed using PolyMorph3D. Each microglia channel was processed separately using the Single Image Main Script to obtain optimal binary volumes of all microglia in each image. Lipofuscin and CD68 channels were processed using the Batch Option B script (Hybrid Batch). A detailed explanation of each script can be found in the section below (see **Figure 1** for workflow summary). Shared measurements between Neurolucida*®* 360 and PolyMorph3D (total microglia branch length, branch points, branch endpoints, and quantity of CD68 pockets overlapping with the microglia volume) were directly compared on a per-cell basis using simple linear regression.

**Figure 1:**
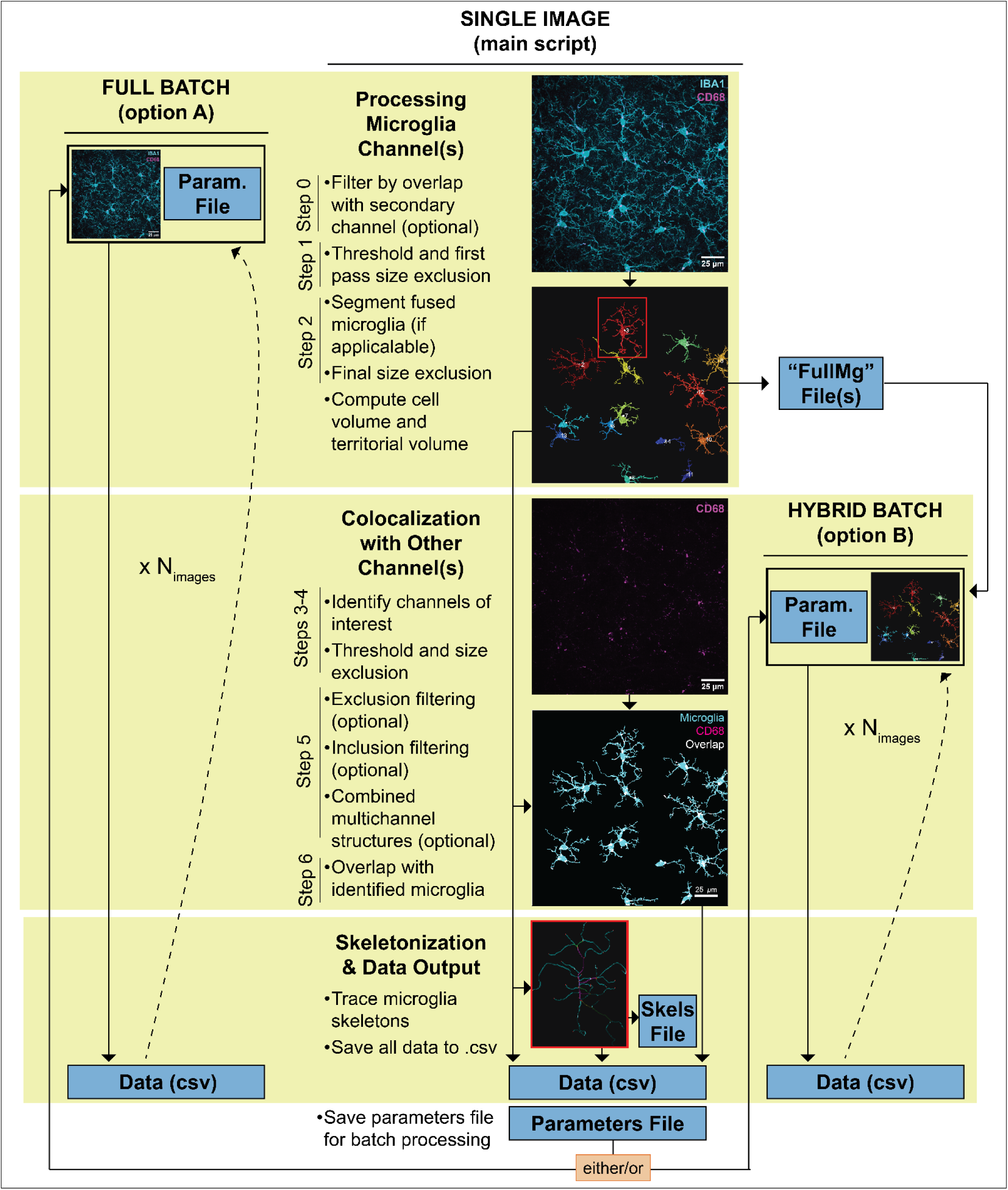
PolyMorph3D workflow overview. The Main Script (Single Image) should be run first. In this script, a single image is manually processed. Individual microglia are identified, cell volume and territory volume of each is calculated, and linear voxel indices for each cell are saved as a .mat file (titled “FullMg”). Users may also use the pipeline to colocalize these microglia volumes with other channels and trace skeletons of each cell for additional morphological data (skeletons will also be saved as .mat file if you wish to reload and change how they appear later). Images can be saved throughout to document progress and will follow the Steps 0-6 shown. After completion, the Main Script saves a Parameters file which can subsequently be loaded into either of the Batch Process scripts. The Full Batch script (Option A) uses the same Parameters for all images and therefore applies the analysis pipeline in a fully automated fashion. The Hybrid Batch script (Option B) requires that any desired images first be run through the Processing Microglia Channel(s) steps of the Main Script (you can skip steps 3-6), which saves identified microglia for each image as “FullMg” .mat file. Thus, microglia thresholding and filtering can be done on a per-image basis while the remaining operations (colocalization, skeletonization) can be done fully in batch. This mode may be useful if you find that a single threshold for all images does not ideally capture microglia structure for all images.

Software Accessibility: All code for PolyMorph3D can be obtained from the following GitHub repository: https://github.com/andrew-yoblinski/PolyMorph3D.git

## RESULTS

PolyMorph3D is a script-based MATLAB program mediated by graphical user interface (GUI) apps; thus, coding experience is not necessary. The user can start with a 3D stack (must be TIFF), threshold and identify valid structures within channels, then colocalize structures from up to 4 channels with the identified microglia. After processing a single image manually with the Main Script, a parameters file is saved and may be used for batch processing many TIFF images (**Figure 1**). Each of these steps is described below.

### Processing Microglia Channel(s)

After selecting the 3D image stack and specifying the primary microglia channel, the user can also specify a secondary microglial channel, which may be used to filter microglia by overlap with a validation marker (ex. Tmem119, DAPI) or merged with the primary channel for improved structural tracing (ex. P2RY12, Cx3Cr1, Tmem119). This option is demonstrated in **Figure 2**, where Iba1 is used as the primary microglia structure channel and the nuclear stain DAPI is used for validation (**Figure 2A-F**). Next, the user thresholds the microglia channel(s) (**Figure 2B-C**). During this step, the user may also specify microglia size range cutoffs. We recommend using a conservative minimum (<50µm^3^) to remove small artifacts that increase processing time; there is another opportunity to further refine these cutoffs in the next steps. We also recommend only using the maximum size cutoff to exclude obvious, excessively large artifacts. The user is encouraged to use the 3D preview options to help set these parameters. After thresholding, the user proceeds through the same cell segmentation and final size cutoff workflow offered in 3DMorph.

**Figure 2:**
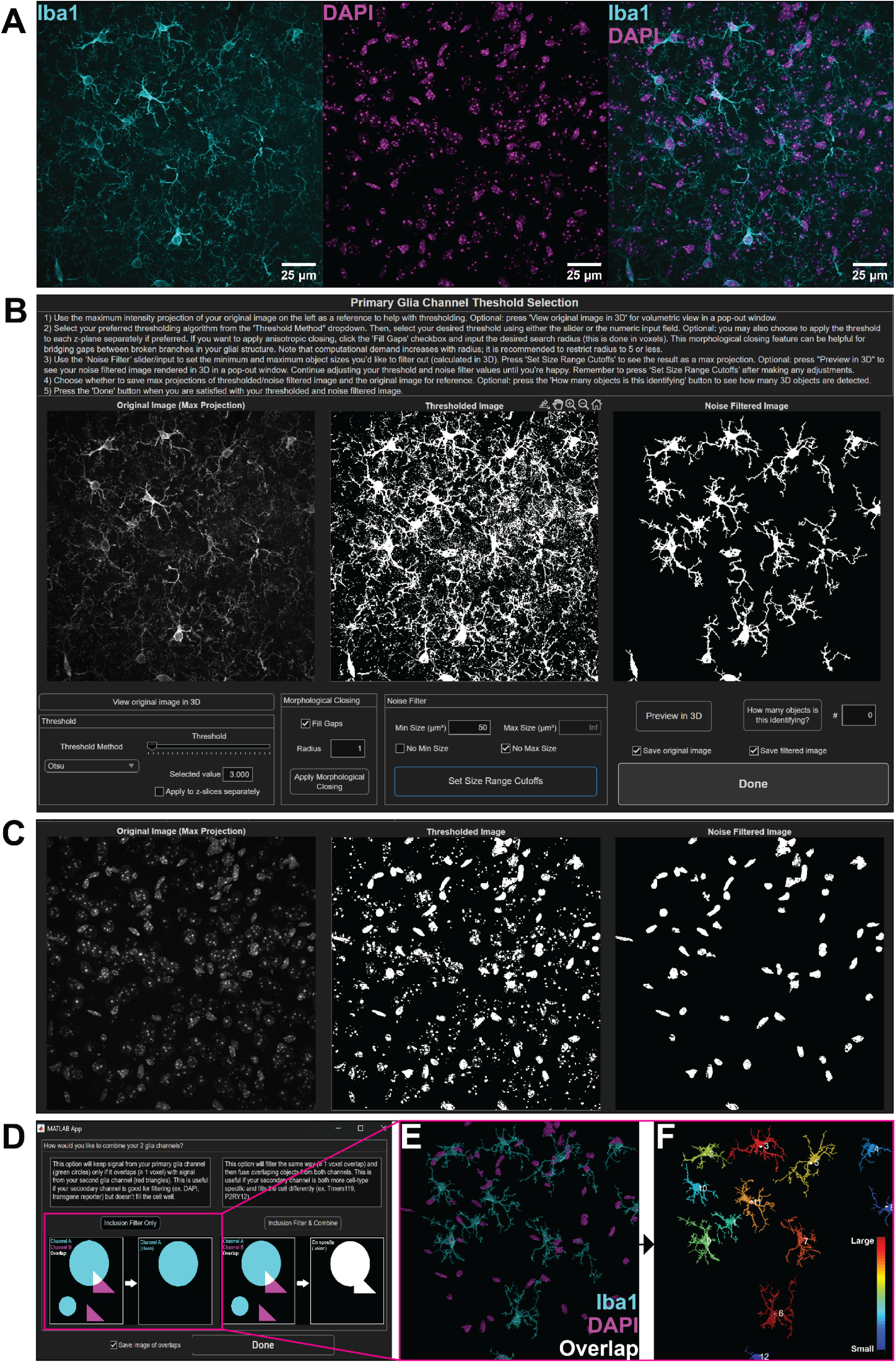
Thresholding and filtering microglia channel(s). **A)** Maximum intensity projections of primary microglia structure channel (Iba1), secondary channel (nuclear stain DAPI) and merge. **B)** Graphical User Interface (GUI) for thresholding and setting initial size range cutoffs for primary microglia channel (Iba1) and **C)** secondary microglia filtering channel (DAPI). **D)** GUI prompting the user to either filter signal from the primary channel (Iba1) based on overlap with secondary channel (DAPI) or filter and combine channels. **E)** Overlap of thresholded primary and secondary channels and **F)** final selection of filtered microglia.

### Colocalization With Other Channel(s)

Before proceeding to the microglia skeletonization step, the user may choose to colocalize the identified microglia traces with up to 4 other channels in the original image stack. If the user is not interested in colocalization, these steps can be skipped.

The user first thresholds each colocalization channel. The user may also set size range cutoffs for any of the colocalization channels, useful for identifying structures expected to be within a narrow size range, such as Shank3 puncta for dendritic spines, and for excluding excessively large structures due to noise, such as lipofuscin (**Figure 3**). After thresholding, there are several options for further signal processing, each intended to meet unique requirements of complex multi-channel designs and can be skipped if not applicable.

**Figure 3:**
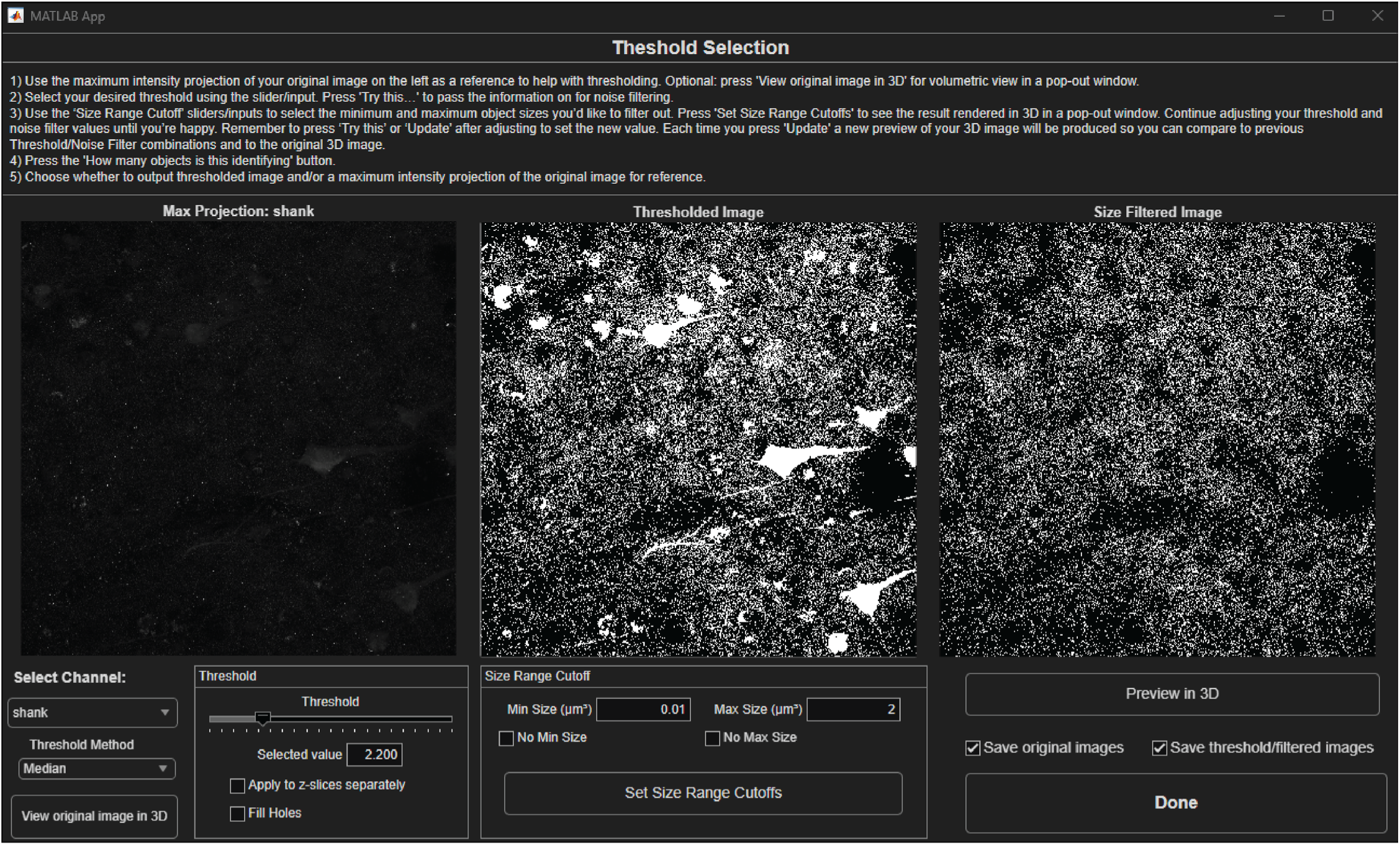
Thresholding and size filtering other channels for colocalization. This figure is a screenshot of the GUI for non-microglia channels. As an example, the thresholding and size exclusion parameters for the Shank3 channel are shown. In the maximum intensity projection (leftmost panel) and the thresholded image (middle panel), one can easily discern the presence of autofluorescence due to lipofuscin accumulated in the cell body. The Size Range Cutoff tool (middle panel) can be used to exclude much of this contaminating signal by restricting identified objects to a plausible range for dendritic spines (rightmost panel).

### Exclusion Filter

The first post-threshold processing step allows the user to filter signal from one channel (A) based on overlap with another channel (B). For instance, in human postmortem brain tissue, accumulation of broadly autofluorescent lipofuscin is well documented (Seehafer & Pearce, 2006; Terman & Brunk, 2004). One method to minimize any confounding effects of lipofuscin is to collect it in its own channel as a comparator (Rocco et al., 2017; Rocco et al., 2016). In PolyMorph3D, if autofluorescence has been collected in Channel B (magenta triangles) and one is only interested in keeping signal from Channel A (cyan circles) that does not overlap with autofluorescence, the user can choose to remove overlapping voxels, remove entire objects, or both (**Figure 4A**). Here, we remove signal from the CD68 channel that overlaps with lipofuscin (**Figure 4B-C**).

**Figure 4:**
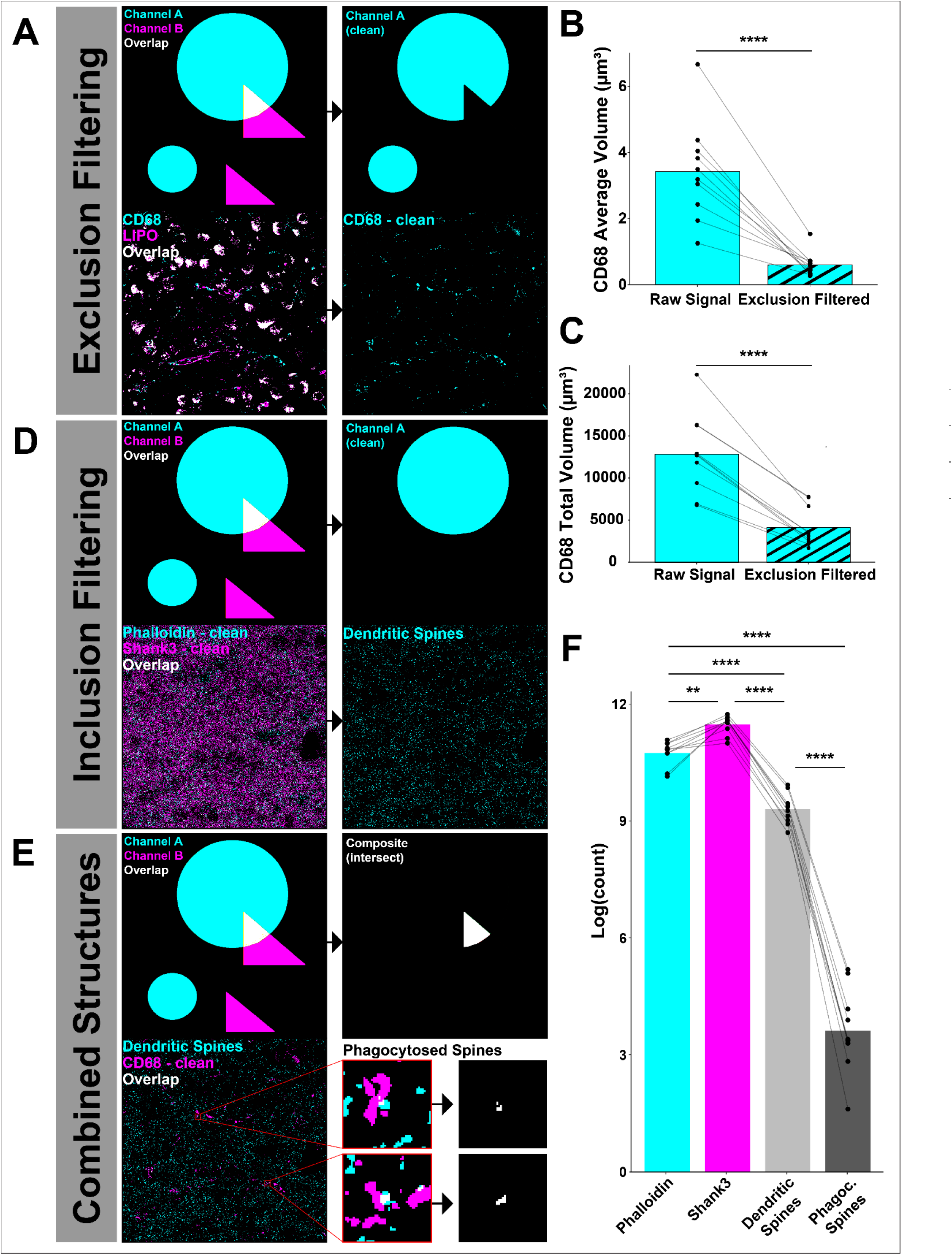
Post-threshold filtering and processing of other channels for colocalization. **A)** Example images of Exclusion Filtering step, with cartoon depicting removal of overlapping voxels between specified channels (upper panel) and real-world example where CD68 signal is filtered to exclude overlap with lipofuscin (lower panel). Validation of clean signal shown quantitatively by (**B**) reduced average volume of identified CD68 objects (p < 0.0001, paired t-test) and (**C**) reduced total volume of CD68 (p < 0.0001, paired t-test) per image after size exclusion filtering. **D)** Example images of Inclusion Filtering step, with cartoon depicting inclusion of overlapping objects between specified channels (upper panel) and real-world example where Phalloidin objects are kept if they overlap with Shank3 (lower panel). Phalloidin fills the volume of a dendritic spine more fully than Shank3 but is not as specific. This filtering procedure allows for reliable identification of complete dendritic spines. **E)** Example images of the Combined Structures filtering step, with cartoon depicting retention of overlapping voxels between specified channels (upper panel) and real-world example where dendritic spine voxels that overlap with CD68 are retained as putative phagocytosed dendritic spines (lower panel). **F)** Quantification of identified phalloidin, Shank3, dendritic spines, and putative phagocytosed spine objects, log transformed to aid with visualization (p < 0.0001, rmANOVA). As expected, there are fewer verified dendritic spines than phalloidin or Shank3 objects (p < 0.0001, post-hoc), and there are fewer phagocytosed spines than total dendritic spines (p < 0.0001, post-hoc). For all analyses: *p < 0.05; **p < 0.01; ***p < 0.001; ****p< 0.0001.

### Inclusion Filter

You may want to keep signal from one channel (A) only if it overlaps with another channel (B) (**Figure 4D**). Here, we use Shank3 to identify dendritic spines, but this excitatory post-synaptic scaffolding protein does not always fill the entire spine volume. Thus, we also stain with phalloidin, an actin filament toxin which fills the entire spine volume but is not specific to dendritic spines. To get the best of both worlds, this step keeps entire phalloidin puncta only if overlapping with Shank3.

### Composite Structure Creation

Finally, the user may specify combinations of channels to form unique composite structures and/or keep only those voxels that overlap. In the example here, we keep only dendritic spine voxels that overlap with CD68. Thus, we measure the level of phagocytosis of dendritic spines based on overlap with microglia and CD68 (**Figure 4E**). As expected, the abundance of dendritic spines identified by colocalization of 2 markers is lower than when using Shank3 and Phalloidin alone, and the number of putative phagocytosed spines is lower still than the total number of dendritic spines identified (**Figure 4F**). Having processed channels for colocalization, the user can now move to automated skeletonization and data output.

### Validation Against Neurolucida® 360

To demonstrate the workflow fidelity described above, we conducted a validation experiment with an existing dataset in human postmortem brain. A random selection of 40 microglia was hand traced using Neurolucida*®* 360 and analyzed with PolyMorph3D in using the Partial Batch script (**Figure 5A**). Morphology metrics shared between the two approaches were compared on a per-cell basis using simple linear regression. PolyMorph3D performs well on branching morphology metrics, with significant correlation with Neurolucida*®* 360 hand traced measurements for total branch length (**Figure 5B**), number of branch points (**Figure 5C**), and number of branch endpoints (**Figure 5D**). Similarly, PolyMorph3D effectively captures CD68 signal that colocalizes with microglia, yielding results comparable to that of Neurolucida*®* 360 (**Figure 5E**).

**Figure 5:**
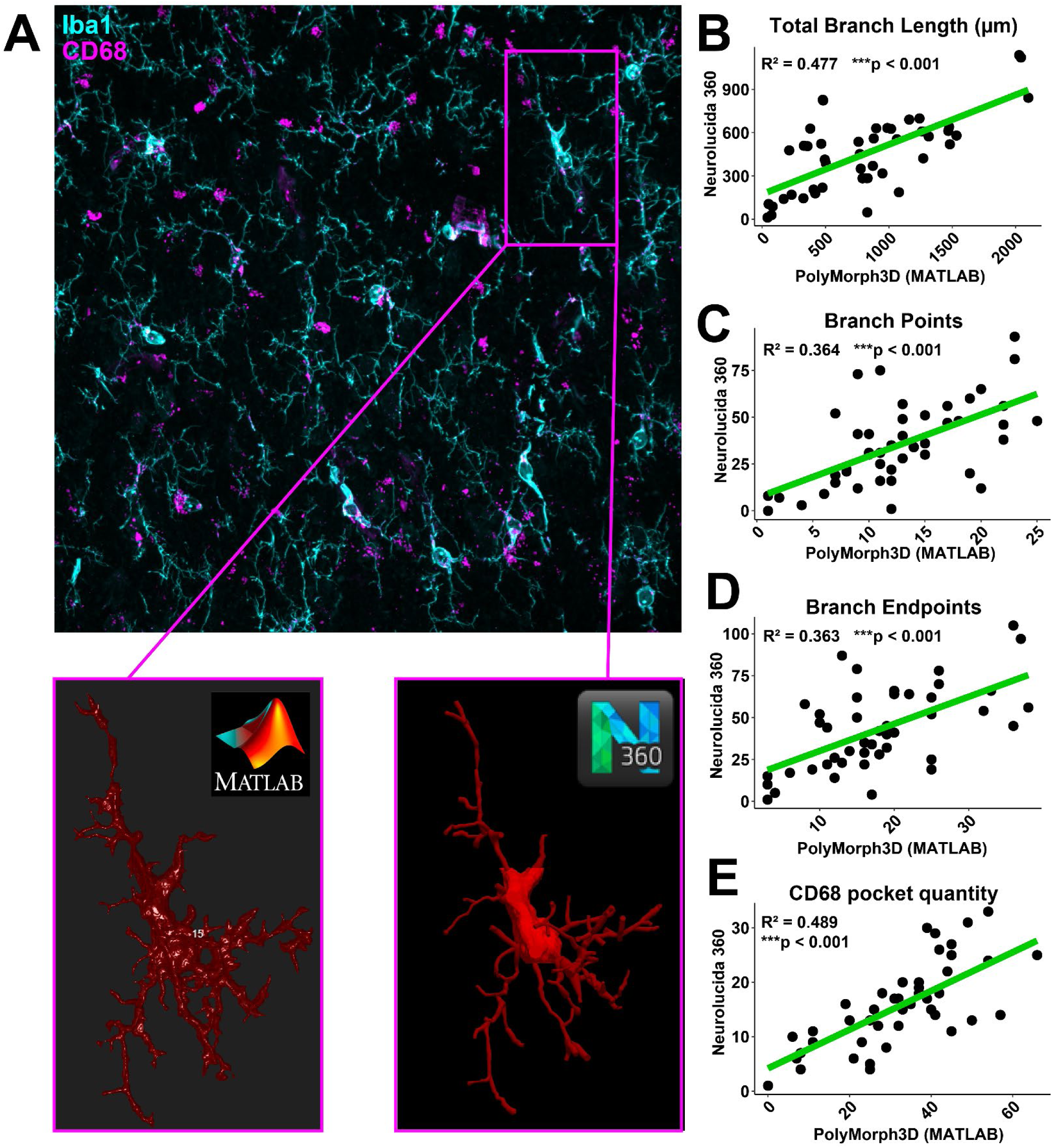
PolyMorph3D captures cell-level information with performance comparable to that of Neurolucida*®* 360. **A)** Example image from human postmortem brain stained for Iba1, CD68, Phalloidin, and Shank3 (upper panel). A random selection of 40 microglia was chosen for validation. Microglia were identified by Iba1 signal using PolyMorph3D (lower left panel) and semi-automatically traced in N Neurolucida*®* 360 (lower right panel). **B)** Total branching length for each cell as determined by PolyMorph3D is significantly correlated (p < 0.001; R^2^ = 0.477) with the Neurolucida*®* 360 value for the same measurement. **C)** Number of branch points for each cell as determined by PolyMorph3D is significantly correlated (p < 0.001; R^2^ = 0.364) with the Neurolucida*®* 360 value for the same measurement. **D)** Number of branch endpoints for each cell as determined by PolyMorph3D is significantly correlated (p < 0.001; R^2^ = 0.363) with the Neurolucida*®* 360 value for the same measurement. **E)** Number of CD68 pockets colocalized with each cell as determined by PolyMorph3D is significantly correlated (p < 0.001; R^2^ = .489) with the Neurolucida*®* 360 value for the same measurement. For all analyses: *p < 0.05; **p < 0.01; ***p < 0.001; ****p< 0.0001.

### Batch Processing Options

After running the Main Script, a .mat file containing batch processing parameters is saved (“Parameters_ImageName.mat”) along with a .mat file containing the voxel indices of identified microglia (“FullMg_ImageName.mat”). The user can now batch process multiple images. There are two options for batch processing (**Figure 6**). The Batch Process Option A (Full Batch) script enables complete analysis in a fully automated fashion. In our preliminary studies, we found that faithfully identifying the full microglia arbor across multiple images is challenging with a single threshold. Thus, we designed Batch Process Option B (Hybrid Batch) to allow the user to identify microglia using the Single Image script for each image, then use the Option B script for remaining steps. Thus, microglia thresholding and filtering can be done on a per-image basis while the remaining operations (colocalization, skeletonization) done fully in batch.

**Figure 6:**
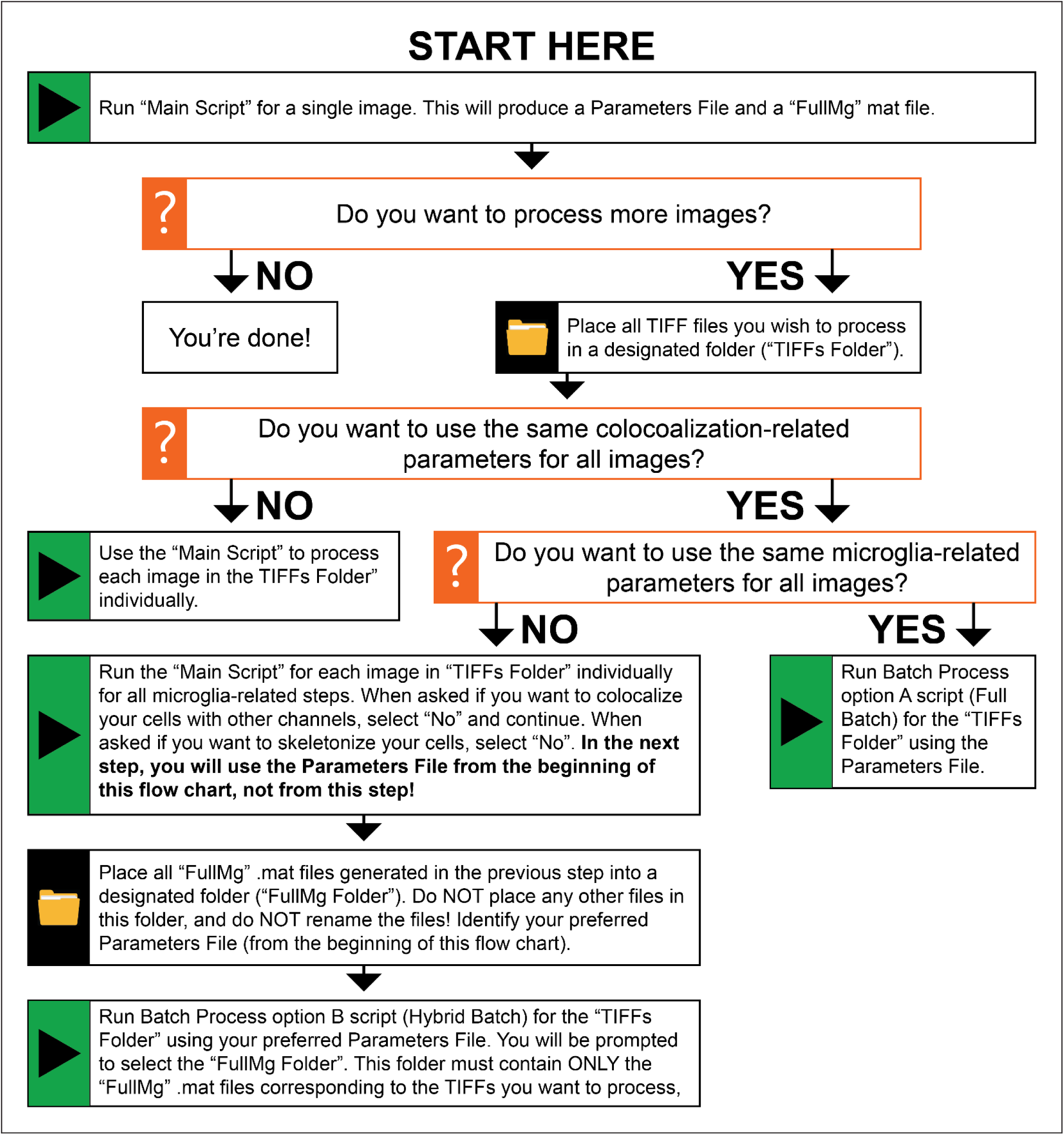
Flowchart for how to use PolyMorph3D. In all cases, you must start with the Main Script and run it from start to finish. This produces a parameters file which stores all of your processing choices to be used for batch processing if desired. There are two options for batch processing. Batch Process Option A script allows the user to analyze images as specified in the Parameters file in a fully automated fashion for any number of images. Alternatively, the Batch Process Option B (Hybrid Batch) script may be used if you want to specify unique parameters for the microglia in each image, while using the same parameters for all colocalization steps. In this script, the user must identify microglia in the Single Image script for each image first and move all FullMg files to a separate folder before proceeding to the Option B script. In the Option B script, the user will load the Parameters file, the FullMg folder, and TIFF image folder. It is critical that you keep the FullMg and TIFF filenames as they are; do NOT change either of them as the script looks for a hard-coded naming pattern between the file types to match them exactly. Thus, microglia thresholding and filtering can be done on a per-image basis while the remaining operations (colocalization, skeletonization) can be done fully in batch. If you want to process multiple images and specify unique parameters for every image, you can process them manually one at a time using the Main Script.

## DISCUSSION

PolyMorph3D fills in analytical and practical gaps left by existing programs for investigating microglia structure and function from fluorescence microscopy, making it a one-stop-shop for these datasets. Our program features the following: 1) high-throughput, 2) high-fidelity for 3-dimensional structure, 3) individual cell-level resolution, 4) colocalization with other channels, and 5) accessible (i.e., low/no cost and easy to use without coding experience). Previously published, 3DMorph (York et al., 2018) served as the backbone for the present program, thus PolyMorph3D retains much of the 3DMorph workflow. However, 3DMorph has limited thresholding and preview options, and critically lacks colocalization capability. PolyMorph3D addresses each of these limitations. First, our program adds several thresholding algorithm options (Otsu, Multi-Otsu, Median, Mean) where 3DMorph restricts the user to Otsu only, and adds an anisotropic closing (specifying a search radius of up to 5 pixels with the imclose function) “Fill Gaps” option which can be useful for bridging gaps in broken branches. Similarly, the 3DMorph noise filter is one-sided and restrictive, constraining the user to a minimum size cutoff in stepped size intervals. PolyMorph3D offers improved flexibility by making size cutoff two-sided (a minimum and maximum cutoff) and accept any numeric. Second, 3DMorph shows previews of only the middle slice in a z-stack, which does not give the user a good sense of either the original image or how well the thresholded/filtered image matches. PolyMorph3D uses the more common maximum intensity projection for 2D previews and the native volshow function for 3D previews. The third and the most significant improvement in PolyMorph3D is the capability to process up to 4 additional channels for overlap with identified microglia. This colocalization capability, as discussed previously, makes PolyMorph3D unique amongst open-source microglia analysis tools (Benkeder et al., 2025; Hansen et al., 2022; Heindl et al., 2018; Salamanca et al., 2018). The colocalization settings can be tailored to a variety of complex experimental designs, such as exclusion and inclusion filtering, and composite object creation. While commercially available Neurolucida*®* 360 and Imaris have similar features, the licenses for these programs can be prohibitively expensive. By contrast, an academic license for MATLAB is comparatively inexpensive and often free to researchers through their institutions. Moreover, PolyMorph3D exhibits comparable performance to that of Neurolucida*®* 360, but with only a fraction of the human labor investment. We estimate the Neurolucida*®* 360 tracing in this study took approximately 40 labor hours to complete, compared to 4 labor hours with PolyMorph3D, around a 10-fold reduction.

In summary, we present a MATLAB-based image analysis solution that interrogates microglia 3-dimensional structure and colocalization with up to 4 other channels with high fidelity and at individual cell-level resolution. PolyMorph3D implements these analyses in a high-throughput fashion through batch processing, is flexible to accommodate a wide range of experimental needs, and requires no prior coding experience. We hope that this low-cost tool will be used by fellow microglia researchers to increase analysis efficiency while capturing the nuanced characteristics of these cells that have proven to be central in brain health and disease.

### Technical Considerations

#### Thresholding

The “Fill Holes” option while thresholding colocalization channels is not the same as the “Fill Gaps” option while thresholding microglia channels. Instead, the relevant function is bwmorph3 using the “fill” operation, which fills isolated interior voxels

##### Colocalization

In all cases, colocalization is defined as overlap of at least 1 voxel. This is true even if the user is interested in object colocalization. In other words, even very minimal overlap between two objects (a single voxel) is sufficient to count two objects as overlapping. Thus, it is highly recommended that the user visually double check outputs to ensure the chosen parameters produce believable results. Images of overlap can be saved and will contain “step6” in the file name. These images are rendered with all microglia, but only objects from other channels that overlap with microglia. Conclusions drawn from colocalization can be based on number of objects alone and/or include consideration of volume of overlap.

##### Visualization Colors

Maximum intensity projections of original TIFF images are always rendered in grayscale. The thresholded images are rendered using the “hot” colormap where color temperature corresponds to voxel location in the z-axis, thus rendering an image that appears more 3D-like. When rendering overlap images, the primary microglia channel is rendered in cyan and other channels in magenta, such that overlap is rendered in white. These color choices were made for optimal viewing and have no relation to original channel wavelength or channel order.

##### 2D vs 3D Visualization

Throughout the script, the user is presented with the opportunity to save images visualizing the inputs and outputs of each processing step either in 2D or 3D. Rendering in 3D can be computationally demanding and time consuming, so we recommend saving 2D visualizations for batch processing purposes. However, it is noteworthy that 2D visualization is a maximum intensity projection and therefore susceptible to artifactual rendering of overlap between channels for voxels that are located at the same XY coordinates but in different Z-planes. Please note that this does NOT apply to the data output: data processing is always maintained in 3 dimensions, this is a quirk of 2D visualization only. As such, we recommend using the 3D rendering option for preliminary parameter optimization and for publication quality rendering needs.

##### Data Output

The skeletonization step produces the following microglia morphology information: total branching length, number of branches, number of branch points, and number of endpoints. As is the case with 3DMorph, this step can either trace only major branches or trace all small processes (note: keeping small processes is much more time consuming). The cell volume and territorial volume of each microglia is also saved. In addition to these single-cell resolution outputs, the script produces image-level data for mean fluorescent intensity for all channels processed, total microglia volume, image volume, maximum microglia volume, average distance between microglia, microglia density, total number of objects in other processed channels, density of objects in other processed channels, and total and average volume of objects in other processed channels.

## Acknowledgements

The authors thank their colleagues in the Translational Neuroscience Program at the University of Pittsburgh for their detailed feedback and insights throughout the development and beta testing of the program. To expedite the development process, AI was used to generate some initial code blocks, with all such generated code subsequently validated and debugged by hand.

